# A nonsense mutation in *TFEC* is the likely cause of the recessive piebald phenotype in ball pythons (*Python regius*)

**DOI:** 10.1101/2020.10.30.362970

**Authors:** Alan Garcia-Elfring, Heather L. Roffey, Andrew P. Hendry, Rowan D. H. Barrett

**Author notes:** The corresponding author’s.

## Abstract

Captive-bred ball pythons (*Python regius*) represent a powerful model system for studying the genetic basis of colour variation and Mendelian phenotypes in vertebrates. Although hundreds of Mendelian phenotypes (colour morphs) affecting colouration and patterning have been described for ball pythons, the genes causing these colour morphs remain unknown. Here, we used crowdsourcing of samples from commercial ball python breeders to investigate the genetic basis of a classic phenotype found in the pet trade, the piebald [characterized by dorsolateral patches of unpigmented (white) skin]. We used whole-genome sequencing of pooled samples followed by population genetic methods to delineate the genomic region containing the causal gene. We identified *TFEC* of the MIT-family of transcription factors as a candidate gene. Functional annotation of SNPs identified a nonsense mutation in *TFEC*, which we conclude is the likely causal variant for the piebald phenotype. Our work shows that ball python colour morphs have the potential to be an excellent model system for studying the genetic basis of pigment variation in vertebrates, and highlights how collaborations with commercial breeders can accelerate discoveries.

## Introduction

The study of colour variation has a long history in evolutionary biology and genetics (Caro 2017; Cuthill et al. 2017; Harris et al. 2019). Over 150 years ago, crossing yellow and green peas helped Mendel to discover that traits are inherited as discrete hereditary units or “factors” (Mendel 1866; reviewed by Ellis et al. 2011). Once Mendel’s work was rediscovered in 1900, invertebrate colour variation helped establish that Mendel’s “factors”, which we now call genes, were located on chromosomes (Bridges 1919; Harrison 1920). Further, colour variation in vertebrates has provided insights in many fields, particularly the biomedical sciences (e.g. Pingault et al. 1998; Tachibana et al. 2003; Richards et al. 2019), developmental biology (e.g. Cohen et al. 2016; Haupaix et al. 2018), and evolutionary biology (e.g. Schweizer et al. 2018; Barrett et al. 2019; Burgon et al. 2020; reviewed by San-Jose and Roulin 2017). However, our current knowledge of the genetics of vertebrate colouration comes primarily from studies on model species, particularly the mouse (e.g. Baxter et al. 2004; Hoekstra 2006; San-Jose and Roulin 2017; but see Ullate-Agote et al. 2020). This bias in the literature provides a limited view of the ways in which genetic variation can affect vertebrate colouration.

Although mice, as all mammals and birds, have one pigment-producing cell type (melanocytes) which synthesizes melanin, reptiles, like fish and amphibians, have three colour-producing cells or chromatophores (reviewed by Bechtel 1978; Mills and Patterson 2009; Olsson et al. 2013). In addition to melanin-producing cells, called melanophores, reptiles have xanthophores and iridiophores that contribute to colouration. Xanthphores synthesize yellow pteridine-based pigments and can also contain red carotenoid pigments that are obtained from diet (often called erythrophores). Iridiophores do not contain pigments but instead produce structural colouration by reflecting light on guanine nanocrystals (Teyssier et al. 2015). The three colour-producing cells interact in 3D space (Grether et al. 2004; Saenko et al. 2013) to generate the remarkable range of colour variation observed in reptiles.

We propose that the ball python (*Python regius*; Figure 1a) is an excellent model for studying the genetic basis of vertebrate colouration (Irizarry and Bryden 2016). Ball pythons are native to western Sub-Saharan Africa, with a range extending from Senegal to Uganda. With a maximum length of approximately 6 feet long, ball pythons are small relative to their better-known Asian congener, the Burmese python (*Python bivittatus*). Owing to their small size, docility, and simple husbandry, ball pythons rose in popularity in the pet trade, particularly among reptile enthusiasts. An additional factor that has contributed to the ball python’s popularity was the discovery of rare phenotypes that affect colouration. In Ghana, claims of rare wild ball pythons with altered patterning go back as early as 1966. In the early 1990s, one such ball python was caught, a piebald. This individual had patches of unpigmented (white) skin (e.g. Figure 1b-d), a pigment deficiency found in many vertebrates (Ahi and Sefc 2017), including humans (Oiso et al. 2013). However, it would take a few years to rule out environmental factors as a causal explanation for the piebald phenotype in the ball python caught from the wild. By 1998, Peter Kahl had reproduced the piebald phenotype in captivity (called ‘proving’ by commercial breeders) and showed it had a recessive mode of inheritance (*A brief history of Royal Ball Python morphs for beginners*, n.d.). This pioneering work of breeders and hobbyists spurred the discovery and propagation of numerous additional and highly diverse colour morphs (e.g. Figure 1b-g). Today, 314 Mendelian phenotypes (i.e., phenotypes that segregate as a single Mendelian factor or major quantitative trait locus) are bred in captivity (known as ‘basic’ morphs; http://www.worldofballpythons.com/morphs), providing a valuable resource for studying the genetics of vertebrate colouration in great detail. Indeed, commercial breeders have taken advantage of epistasis and produced over 6000 unique phenotypes (‘designer morphs’) with different combinations of the Mendelian colour factors.

**Figure 1.**
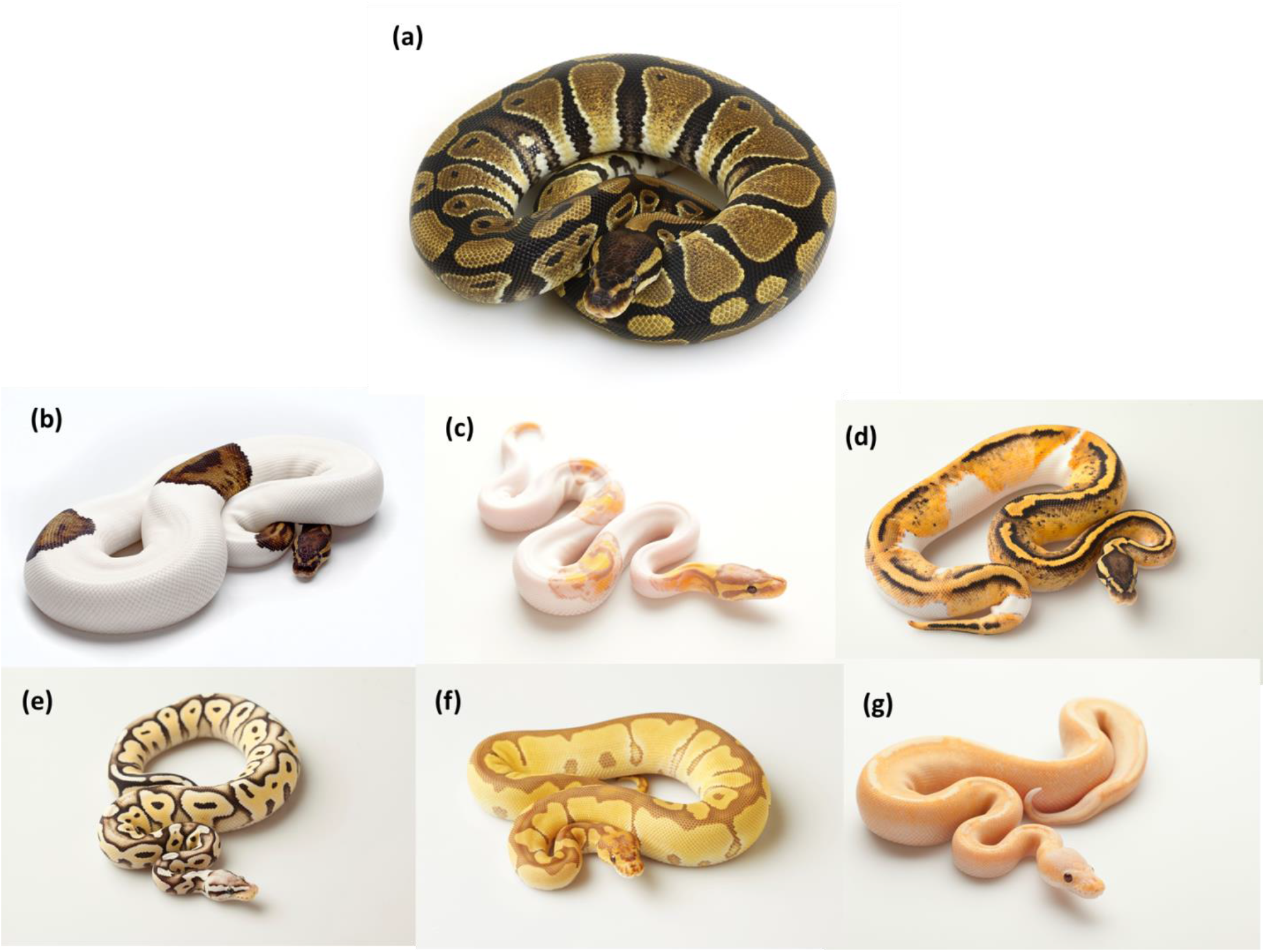
A small sample of the phenotypic variation found in captive-bred ball pythons (*Python regius*). (a) wild-type, (b) piebald, (c) banana piebald, (d) pastel piebald, (e) pastel HRA enhancer, (f) ultramel clown, (g) banana champagne. Photo credit (b-g): Designing Morphs.

Ball pythons therefore provide an outstanding opportunity to study the genetic basis of Mendelian phenotypes related to pigmentation and patterning. For example, ball python colour morphs can be characterized by changes to pattern or to pigmentation or both. Phenotypes with changes to pigmentation but not pattern (Figure 1c) are presumably determined by changes to genes with roles in a pigment synthesis pathway. On the other hand, aberrations to pattern, like white-spotting, are thought to occur through mutations in genes with functions in progenitor cell migration and differentiation from the neural crest (reviewed by Mills and Patterson 2009). However, most research on colouration, vertebrate and invertebrate, has focused on pigmentation, particularly melanin-based pigmentation (e.g. San-Jose and Roulin 2018), rather than on patterning (Mills and Patterson 2009). The extensive phenotypic variation in pigment and pattern found in captive-bred ball pythons can therefore help correct the bias (e.g. Ullate-Agote et al. 2020) associated with mammalian models and melanin-based pigmentation in the scientific literature on vertebrate colouration.

In this study, our goal was to discover the candidate mutation(s) that produce the Mendelian phenotype affecting pattern in ball pythons, the piebald (Figure 1b). We used crowdsourcing of samples from commercial breeders, thereby bridging a gap between academia and industry. Genes known to cause white-spotting in other vertebrates, like *KIT, MITF*, *EDNRB*, and *SOX10* (Fleischman et al. 1991; Baxter et al. 2004; Ahi and Sefc 2017) represent *a priori* candidate genes, but given the differences in pigmentation development between reptiles and previously studied model systems, we employed a whole-genome approach to facilitate discovery of unknown candidate genes. We sequenced whole-genomes of pooled samples and then applied population genetic methods to delineate the genomic region of interest and identify candidate genes. We then functionally annotated SNPs to identify the putative causal mutation. Given the character of the piebald phenotype (i.e., dorsolateral patches of unpigmented skin), we expected the candidate gene to be expressed in the neural crest and associated with chromatophore migration and differentiation.

## Methods

### Sampling and DNA extraction

We obtained ball python samples by crowd-sourcing from commercial breeders in Canada (Mutation Creation, T. Dot Exotics, The Ball Room Canada, Designing Morphs), who supplied us with shed skin. We then used shed skin samples for 47 piebald individuals (i.e., individuals homozygous for the piebald variant; Table S1) and 52 non-piebald individuals (Table S2). Although individuals from both sets of samples contained additional basic morphs, the only consistent difference between the two was the piebald phenotype. We attempted to maximize the number of individuals that come from different families to minimize the effects of population structure, although there were some exceptions (Table S2). From each sample we used approximately 0.1 g of shed skin, cut to small pieces using scissors, for DNA extraction. We extracted DNA following a standard phenol-chloroform procedure, with a 24-hour proteinase-K incubation time at 37 °C. Piebald and non-piebald samples were prepared on different working days to avoid contamination. We quantified all samples using a Picogreen® ds DNA assay (Thermo Fisher Scientific, Waltham, USA) on an Infinite® 200 Nanoquant (Tecan Group Ltd. Männedorf, Switzerland).

### Sequencing

After DNA extraction, we mixed DNA of individuals (according to phenotype) in equimolar amounts to obtain a single pool for each phenotype, ‘piebald’ and ‘non-piebald.’ We used PCR-based whole-genome libraries for both pools, which were prepared at the McGill University and Genome Quebec Innovation Center, Montreal, Canada. We sequenced 150 bp paired-end reads using two lanes of Illumina HiSeqX.

### Bioinformatics

To process raw reads, we applied filters based on read quality and length, keeping reads with a minimum quality of 20 (--quality-threshold 20) and a length of 50 bp (--min-length 50). We then aligned processed reads to the Burmese python (*Python bivittatus*) draft assembly Pmo2.0 (Castoe et al. 2013) using the program *NextGenMap* (Sedlazeck et al. 2013). *NextGenMap* was designed for aligning reads to highly polymorphic genomes or genomes of closely related species. We used *SAMtools* (Li et al. 2009) to convert SAM files to BAM format and remove reads with mapping quality below 20 (samtools view -q 20). We filtered for PCR duplicates using the program *MarkDuplicates* of Picard Tools (Wysoker et al. 2013). We then created a mpileup file (samtools mpilep -B) from which the synchronized (sync) file was produced using *Popoolation2* (Kofler et al. 2011b). The sync file contains read counts for all nucleotides sequenced in the genome and it is used for subsequent downstream analyses (e.g. F_ST_ scan). We also applied the same protocol as above but instead aligned reads to the chromosome-length Burmese python reference genome, Python_molurus_bivittatus-5.0.2_HiC.assembly (Dudchenko et al. 2017, 2018).

### Identification of fixed SNPs and delineation of the candidate genomic region

We applied an F_ST_ scan to identify fixed SNPs across the two pools. For this procedure, we used the *fst-sliding.pl* script of *Popoolation2* (--min-count 10, --min-coverage 20, --max-coverage 500, --min-covered-fraction 0, --window-size 1, --step-size 1, --pool-size 47:52, --suppress-noninformative). We then identified candidate SNPs as those with F_ST_ estimates of 1 and mapped them to genes. We used a custom BASH script to map fixed SNPs to genes in the gene annotation file using scaffold and SNP position. Because the draft assembly is highly fragmented (Castoe et al. 2013), we also applied the same F_ST_ scan on data aligned to the chromosome-length genome assembly (Dudchenko et al. 2017, 2018) to get a better delineation of the genomic region of interest within candidate SNPs. However, this latter assembly is not annotated with genetic features – hence necessitating the use of both assemblies.

### Functional annotation of SNPs

We next functionally annotated genome-wide SNPs with the software snpEff (Cingolani et al. 2012) to aid in identifying the putative causal mutation for the piebald phenotype. SnpEff was designed for annotating and predicting the effects of SNPs, such as amino acid changes. This program provides an assessment of the impact of a SNP, including ‘HIGH’ (e.g. stop codon), ‘MODERATE’ (e.g. non-synonymous change), ‘LOW’ (e.g. synonymous change), or ‘MODIFIER’ (change in an intergenic area).

## Results

### SNP dataset

From the BAM files, we obtained an average depth of coverage of 50.5 and 52.6 for the piebald and non-piebald pools, respectively. After filtering, our dataset consisted of 3,095,304 SNPs for the draft assembly and 3,221,285 SNPs in the chromosome-length assembly.

### FST scan for fixed SNPs and candidate genomic region

Across all SNPs, we found an average F_ST_ of 0.03456 – relatively low, indicating that population structure was minimized. Using the draft assembly, we identified 129 fixed SNPs (F_ST_ = 1.0) and 369 SNPs with F_ST_ > 0.9. Plotting the F_ST_ values (Figure S1) shows peaks across the reference genome, demonstrating the fragmented state of the draft assembly. In the chromosome-length assembly, we found 131 fixed SNPs (F_ST_ = 1.0) and 372 SNPs with F_ST_ > 0.9. The chromosome-length assembly also shows that 128/131 fixed SNPs and 365/372 SNPs with F_ST_ > 0.9 map to a single region on scaffold 7, clearly delineating a genomic region of interest (Figure 2). The 128 fixed SNPs map to a region 8 Mb long (scaffold 7: 49526089-57612101). This region becomes 26.5 Mb when considering the 365 SNPs with F_ST_ > 0.9 (scaffold 7: 49246858-76754467). However, most of these 365 SNPs cluster within the 8 Mb region demarcated by fixed SNPs, with only 11 mapping beyond the bounds of this region (three upstream, eight downstream).

**Figure 2.**
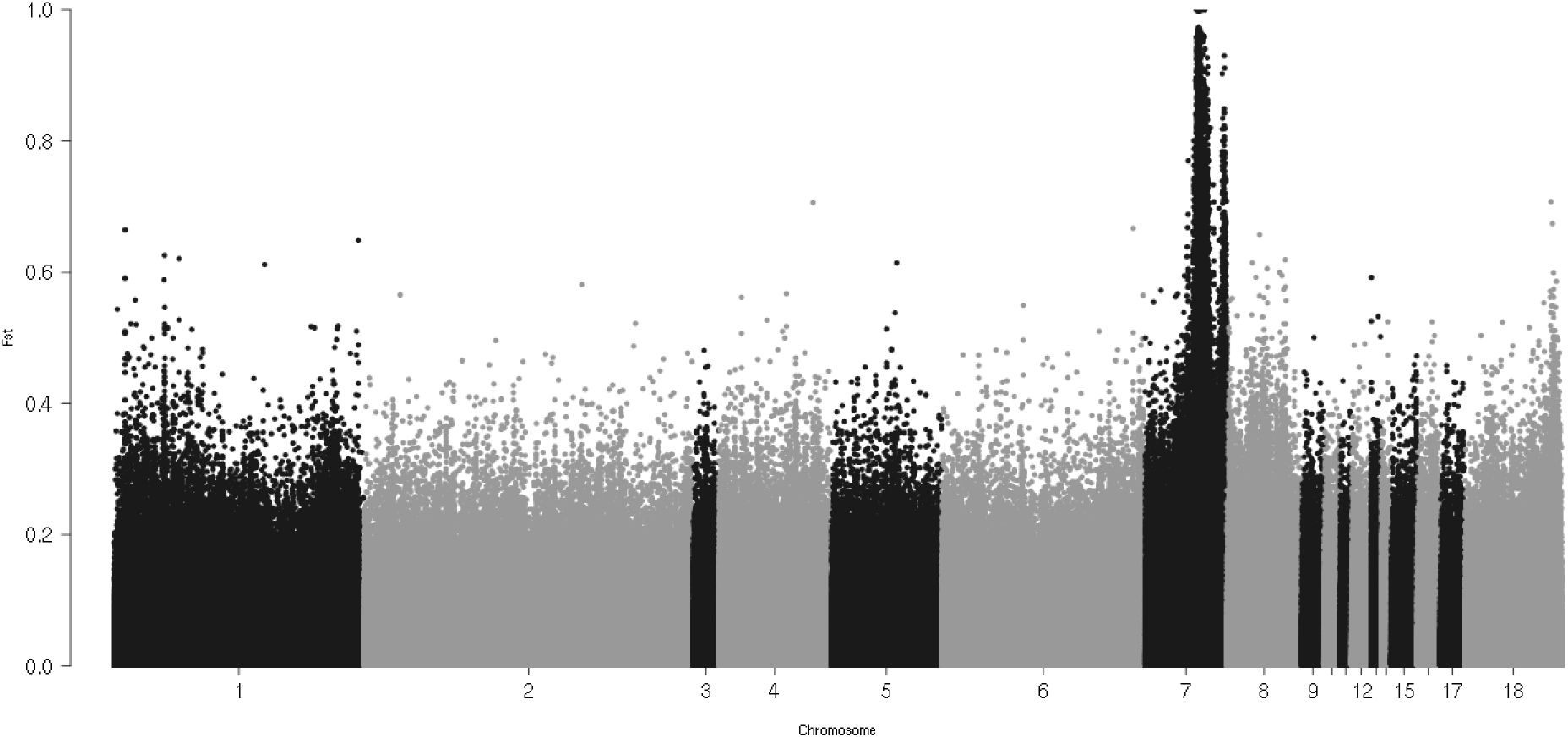
Fst plot between piebald and non-piebald samples on the chromosome-length assembly (Python_molurus_bivittatus-5.0.2_HiC.assembly).

### Candidate genes and SNP annotation

We found that 59 fixed SNPs map to 12 different protein-coding genes (*BMT2*, *CAPZA2*, *CNTN1*, *DOCK4*, *FOXP2*, *GPR85*, *KCND2*, *LOC103067393*, *LSMEM1*, *ST7*, *TES*, and *TFEC*). We annotated SNPs for predicted impact (see methods) and found that all fixed SNPs have the lowest impact classification (MODIFIER), often assigned to mutations in non-functional regions, and most map to introns. However, we found other impact classifications among SNPs with F_ST_ greater than 0.9. One synonymous (i.e., silent) SNP (LOW impact, NW_006539084.1 26867, F_ST_ = 0.92) is located within the gene *LSMEM1* and one nonsense SNP (HIGH impact, NW_006534020.1 160458, F_ST_ = 0.96) is located within the gene *TFEC*. This nonsense variant consisted of a C>T mutation on Arg165, resulting in a premature Opal stop codon (Table S3).

## Discussion

We regard the ball python as a powerful new model for understanding the genetic basis of pigmentation in vertebrates. As proof-of-principle, we set out to discover candidate genes and causal mutations for the classic colour morph found in the pet trade, the piebald. Over the last thirty years, ball python breeders have discovered and successfully propagated in captivity more than three hundred Mendelian phenotypes. Yet, researchers working on the genetics of vertebrate colouration have been slow to appreciate the ball python as a powerful new model (but see Irizarry and Bryden 2016). We set out to bridge this gap. To do so, we appealed to commercial ball python breeders to join our efforts in discovery, starting with uncovering the genetic basis of the piebald phenotype. Commercial breeders have a vested interest in acquiring the ability to identify heterozygotes for recessive phenotypes (i.e., carriers of recessive alleles affecting colouration), and piebald is one such phenotype, although subtle phenotypic differences on heterozygotes are not uncommon (e.g. ring of unpigmented skin, often near the vent).

To elucidate the genetic basis of the piebald phenotype, we combined a case-control experimental design with whole-genome sequencing, population genetic methods, and functional annotation of the candidate SNPs. We identified a premature stop codon in the protein-coding region of the gene *TFEC* as the putative genetic basis for this Mendelian phenotype. *TFEC* belongs to a family of transcription factors (*MITF*) known to be expressed in the neural crest and pigment cells. However, functional validation through gene editing techniques, like CRISPR cas9 (Rasys et al. 2019), is required to establish that the candidate nonsense mutation is the causal variant.

With knowledge of the specific genetic variants underlying pigmentation variation in ball pythons, gene editing could pave the way to produce colour morphs free of secondary phenotypes that are undesired from a commercial or ethical perspective. Genes that contribute to colouration are also expressed in many other tissues derived from neural crest cells (Donoghue et al. 2008). Thus, mutations that alter colour or patterning often alter other traits, such as behaviour (Ducrest et al. 2008; McKinnon and Pierotti 2010). Not surprisingly, there are ball python colour morphs linked to morphological deformities and neurological dysfunction (Rose and Williams 2014). For example, the incomplete dominant ‘spider’ morph is associated with a neurological syndrome called the ‘head wobble’ (Fox and Hogan 2020). The propagation of colour morphs with correlated traits have implications for the health and welfare of ball pythons in captivity (Rose and Williams 2014) and are a source of current controversy among hobbyists and commercial breeders (Riera 2015). In 2017, the International Herpetological Society in the U. K. banned ball pythons with the spider mutation, which is thought to be homozygous lethal, from their reptile shows (International Herpetological Society 2020). However, it is not known if the correlated trait(s) for any particular colour morph is due to pleiotropic effects or linked variation. If due to linked variation, targeted mutation on a different genetic background (i.e., without the linked variant) could allow the reproduction of colour phenotypes free of behavioural and morphological abnormalities. If due to pleiotropy, targeted mutation at different nearby loci could achieve the same effect.

The wide variety of ball python phenotypes found in the pet trade offers the opportunity to study how genes control vertebrate pigmentation and pattern formation. Changes to pigmentation are often caused by mutations in genes involved in the pigment synthesis pathway (Mills and Patterson 2009; Hubbard et al. 2010). For example, albinism occurs when an individual inherits two copies of a gene involved in melanin synthesis (e.g. OCA2) that is non-functional. Snakes with two non-functional OCA2 genes lack dark melanin-based pigment though contain yellow pigments produced in the xanthophores (Saenko et al. 2015). Changes to pattern, on the other hand, are often attributed to genes with functions in cell migration from the neural crest and differentiation into colour-producing cells (Mills and Patterson 2009). However, Mendelian phenotypes characterized by changes to either pigmentation or patterning are likely to map to the protein-coding regions of genes. Genomic research on the most extensively studied vertebrate, humans, shows that Mendelian phenotypes are generally caused by mutations arising in protein-coding regions and affecting protein function (Cooper et al. 2010; Chong et al. 2015). The progenitor cells of chromatophores originate in the neural crest and mutations to these genes result in partial or total loss of pigment and structural colouration. Mutations to genes such as *MITF*, *KIT*, *EDNRB*, and *SOX10*, which are expressed in the neural crest, results in white spotting or a complete lack of pigment across a wide range of vertebrates, including fish, amphibians, birds, and mammals (Baxter et al. 2004; Kelsh 2006; George et al. 2016; Ahi and Sefc 2017; Woodcock et al. 2017; Camargo-Sosa et al. 2019; Goding and Arnheiter 2019). These genes represented *a priori* candidates for the piebald phenotype in ball pythons, but did not show differentiation between our phenotypic pools.

We identified a putative causal mutation for the piebald phenotype in ball pythons located in a new candidate gene for vertebrate pigmentation: the gene *TFEC* (Figure 3, Table S3). This variant, which produces a premature stop codon, was nearly fixed (F_ST_ = 0.96). Complete differentiation between the piebald and non-piebald pools was prevented by a single read for the reference allele that was sequenced in the piebald samples. It is possible this is the result of a sequencing error or the inclusion of an unknown heterozygote. *TFEC* was not one of the candidate genes expected *a priori*. This particular gene has not been implicated in mammalian colouration. However, *TFEC* is part of the MIT-family of transcription factors known to be involved in animal patterning. For example, this family consists of the gene *MITF*, which forms heterodimers with the other three members, *TFEB*, *TFE3*, and *TFEC* (Goding and Arnheiter 2019). In mouse models, double knockout mutations to *MITF* and *TFE3* cause all-white phenotypes (Steingrímsson et al. 2002). Interestingly, the same study showed that knocking out *TFEC* did not result in a phenotype different from the wild-type mouse. There is evidence that shows *TFEC* is expressed in the retinal pigment epithelium of mammals (Rowan et al. 2004). No evidence suggests, however, that *TFEC* is expressed in cells that descend from the mammalian neural crest, which includes pigment-producing cells. In contrast, recent evidence from the zebrafish model shows that *TFEC* is highly expressed in neural crest cells, specifically progenitor pigment cells and iridophores (Petratou et al. 2019; Saunders et al. 2019). In fact, *TFEC* is expressed in two dorsolateral patches on the torso during embryogenesis (Lister et al. 2011; Petratou et al. 2018; Petratou et al. 2019), reminiscent of where the white-spotting occurs in the piebald phenotype in ball pythons. The expression patterns of *TFEC* in zebrafish, together with our results, makes a strong case that *TFEC* is the piebald gene in ball pythons and that a premature stop codon is the likely causal mutation.

**Figure 3.**
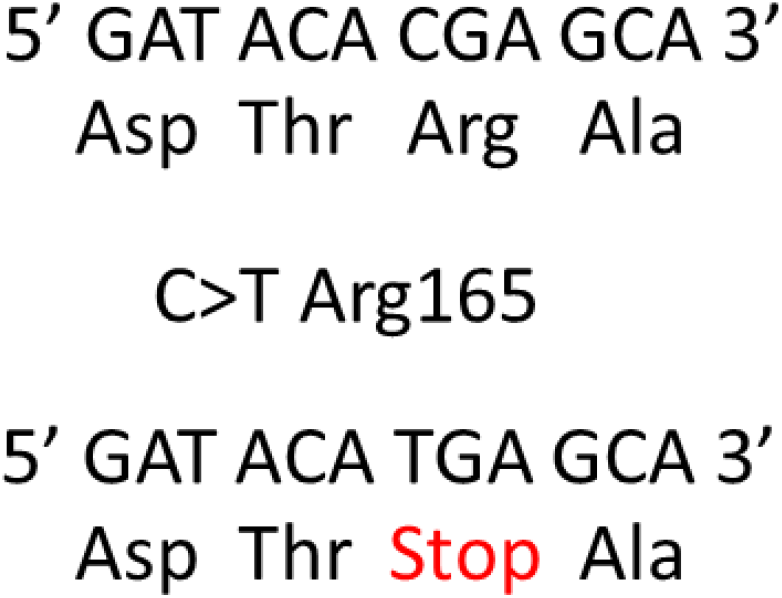
A premature stop codon on Arg165 is the putative causal variant for the piebald phenotype.

## Conclusion

In this study, we aimed to uncover, for the first time, the genetic basis of a ball python colour morph. We discovered that a premature stop codon in the gene *TFEC* is the likely cause for the Mendelian piebald phenotype found in ball pythons from the pet trade. The lack of evidence for *TFEC* affecting mammalian pigmentation highlights the need to study the genetic basis of colouration in non-mammalian vertebrate models. Future work will be aimed at demonstrating causality through direct functional manipulation. These efforts would produce the first genetically engineered ball python, a step that would open the door to safe reproduction of other colour morphs through targeted mutation.

## Acknowledgments

RDHB was supported by an NSERC Discovery Grant and Canada Research Chair. We thank Mutation Creation, T Dot Exotics, The Ball Room Canada, and Designing Morphs for supplying samples.

## Data accessibility

The data that support the findings of this manuscript will be uploaded to Dryad.

## Contributions

AGE, HR, APH, and RDHB conceived and designed the study. HR collected samples. AGE performed the molecular work. AGE performed the bioinformatics and analyzed the genomic data. AGE wrote the manuscript with input from all authors.

## Supplementary figures

**Figure S1.**
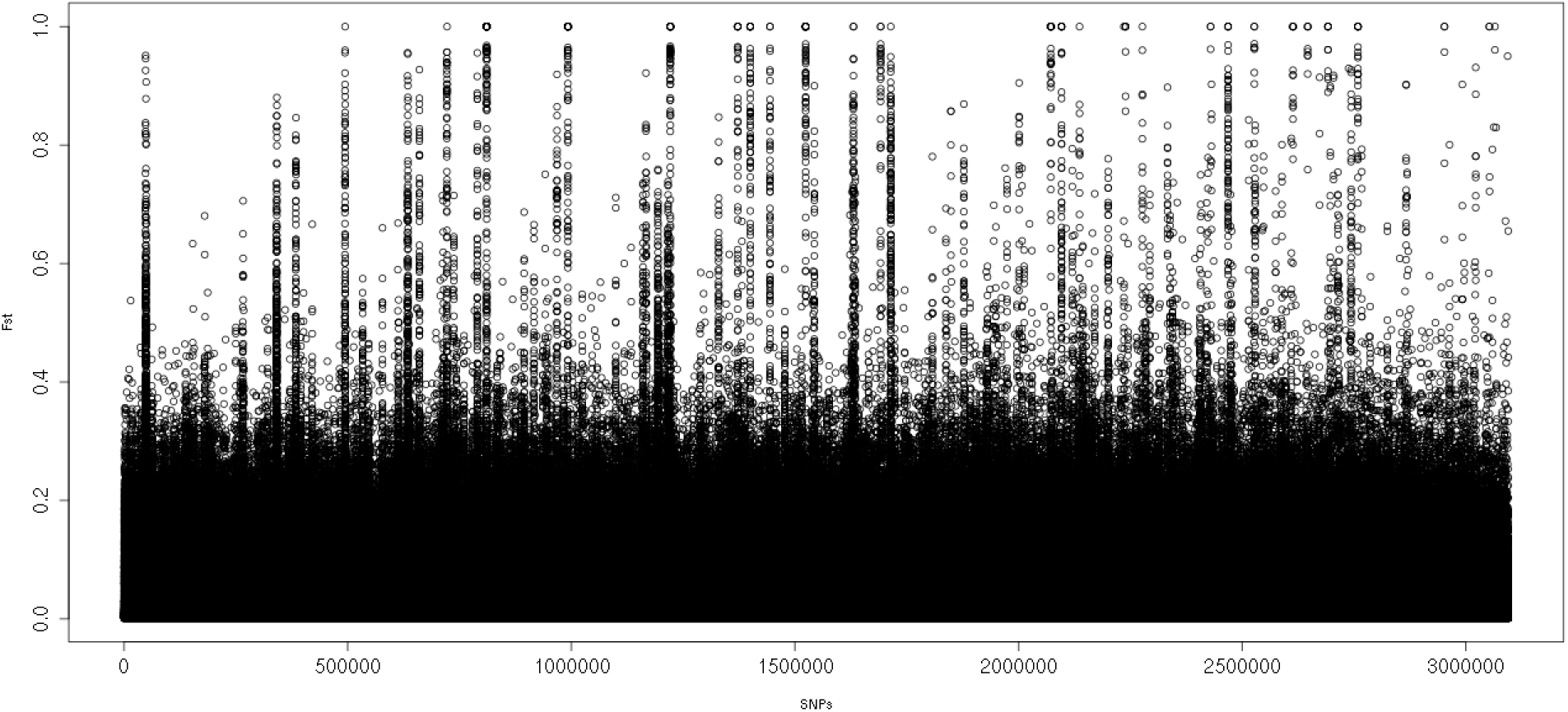
Fst plot of SNPs mapped to the Burmese python draft assembly (Pmo2.0). The 129 fixed SNPs in this plot were used to map to candidate genes.

