## Supplemental Tables 1 and 2 for "A nonsense mutation in *TFEC* is the likely cause of the recessive piebald phenotype in ball pythons (*Python regius*)"

Table S1. Piebald sample ID and additional morphs in each sample. Note that all samples have piebald phenotype. For a description of the different morphs found in the pet trade see http://www.worldofballpythons.com/morphs.

| Sample ID | Morph(1) | Morph(2) | Morph(3) |
|---|---|---|---|
| HR0024 | Piebald | Black-Pastel | |
| HR0028 | Piebald | | |
| MC0023 | Piebald | | |
| MC0039 | Piebald | Fire | |
| MC0075 | Piebald | PVI-Axanthic | |
| MC0082 | Piebald | Pastel | |
| MC0120 | Piebald | Yellowbelly | Het-Lavender |
| MC0124 | Piebald | Super-Pastel | Het-Clown |
| MC0126 | Piebald | Pastel | Fire |
| MC0144 | Piebald | Vanilla | |
| MC0172 | Piebald | | |
| MC0175 | Piebald | Pastel | Yellowbelly |
| MC0196 | Piebald | Pastel | |
| MC0206 | Piebald | Het-Lavender | Albino |
| MC0207 | Piebald | Pastel | |
| MC0222 | Piebald | | |
| MC0269 | Piebald | | |
| MC0284 | Piebald | | |
| MC0299 | Piebald | | |
| MC0305 | Piebald | | |
| MC0307 | Piebald | Albino | |
| MC0314 | Piebald | MJ-Axanthic | |
| MC0336 | Piebald | | |

| | | | |
|---|---|---|---|
| MC0337 | Piebald | MJ-Axanthic | |
| MC0342 | Piebald | | |
| MC0350 | Piebald | | |
| MC0380 | Piebald | Mojave | Banana |
| MC0388 | Piebald | Pinstripe | |
| DM0007 | Piebald | Super-Pastel | |
| DM0009 | Piebald | Black-Pastel | |
| TDOT034 | Piebald | Pastel | |
| TDOT036 | Piebald | | |
| MC0390 | Piebald | Pastel | Yellowbelly |
| MC0391 | Piebald | Orange-Dream | |
| MC0392 | Piebald | Special | |
| MC0393 | Piebald | Orange-Ghost | |
| MC0394 | Piebald | Pastel | Fire |
| MC0395 | Piebald | Pastel | |
| MC0396 | Piebald | | |
| MC0397 | Piebald | Clown | |
| MC0398 | Piebald | | |
| MC0399 | Piebald | | |
| MC0400 | Piebald | Phantom | |
| MC0401 | Piebald | | |
| MC0402 | Piebald | Enchi | |
| MC0403 | Piebald | Yellowbelly | Orange-Dream |
| MC0404 | Piebald | Java | |

Table S2. Non-piebald sample ID and additional morphs in each sample. Note no individuals with the piebald phenotype are included. Individuals with no additional morphs have wild-type colouration. * denotes offspring from dam1 (D1) and sire1 (S1) and ** denotes offspring from dam2 (D2) and sire2 (S2).

| Sample ID | Morph(1) | Morph(2) | Morph(3) | Morph(4) |
|---|---|---|---|---|
| HR0001 | Normal | | | |
| HR0002 | Pastel | Spider | Yellowbelly | |
| HR0003 | Pastel | Specter | | |
| HR0004 | Banana | Pastel | Yellowbelly | |
| HR0005 | Super-Mojave | | | |
| HR0006 (D1) | Black-Pastel | HRA | Pastel | |
| HR0007 (S1) | Cinnamon | | | |
| HR0008 | Champagne | | | |
| HR0009 | Lesser | Yellowbelly | | |
| HR0010 (S2) | Blade | Clown | | |
| HR0011 | Mojave | Special | | |
| HR0012 (D2) | Phantom | | | |
| HR0014 | Black-Pastel | Het-Clown | | |
| HR0015 | Leopard | Spider | | |
| HR0016 | Albino | Enchi | | |
| HR0017 | Blade | Het-Clown | | |
| HR0018 | Black-Pastel | Pinstripe | Het-Albino | |
| HR0019 | Black-Pastel | | | |
| *HR0006-1 | Black-Pastel | Cinnamon | | |
| *HR0006-2 | Black-Pastel | Cinnamon | | |
| *HR0006-3 | HRA | Pastel | | |
| *HR0006-4 | HRA | Pastel | | |
| *HR0006-5 | HRA | Pastel | | |

| | | | | |
|---|---|---|---|---|
| *HR0006-6 | Black-Pastel | HRA | | |
| **HR0012-2 | Phantom | Het-Clown | | |
| **HR0012-3 | Blade | Het-Clown | | |
| **HR0012-4 | Normal | Het-Clown | | |
| **HR0012-5 | Blade | Het-Clown | | |
| MC0001 | Asphalt | Champagne | Enchi | |
| MC0002 | Cinnamon | | | |
| MC0003 | Spider | OD | Yellowbelly | |
| MC0004 | Enhancer | | | |
| MC0005 | Leopard | Pastel | Het-Green Ghost | |
| MC0006 | Mojave | Pastel | Special | |
| MC0007 | Blade | Ghost | | |
| MC0008 | Spotnose | Het-Clown | | |
| MC0009 | Cinnamon | Yellowbelly | Het-Casper Ghost | |
| MC0010 | Cinnamon | Pastel | | |
| MC0011 | Lesser | Goblin | | |
| MC0012 | Het-Clown | | | |
| MC0013 | OD | Super-Pastel | Pinstripe | Spider |
| MC0014 | Banana | Black-Pastel | Pastel | |
| MC0015 | Asphalt | | | |
| MC0016 | Asphalt | Het-Albino | | |
| MC0017 | Enchi | Het-Clown | | |
| MC0020 | | | | |
| MC0021 | Mojave | Spider | | |
| MC0022 | Pastel | | | |
| MC0024 | Super-Asphalt | Enchi | Pastel | |
| MC0026 | Cinnamon | Clown | Pastel | |
| MC0027 | Enhancer | | | |
| MC0028 | Butter | Enchi | | |
