## Supplemental Table 3 for "A nonsense mutation in *TFEC* is the likely cause of the recessive piebald phenotype in ball pythons (*Python regius*)"

Table S3. Annotated SNPs with Fst > 0.9

| Scaffold | Position | Ref. Allele | Alt. Allele | Effect<br>Effect Seq. Ontology | Impact | Gene or genic region |
|---|---|---|---|---|---|---|
| NW_006532038.1 | 193024 | A | G | intron_variant | MODIFIER | CADPS2 |
| NW_006532038.1 | 238074 | C | T | intron_variant | MODIFIER | CADPS2 |
| NW_006532038.1 | 251594 | T | G | intron_variant | MODIFIER | CADPS2 |
| NW_006532038.1 | 715684 | G | A | 3_prime_UTR_variant | MODIFIER | ASB15 |
| NW_006532398.1 | 164921 | G | C | intergenic_region | MODIFIER | KCND2-KCND2 |
| NW_006532398.1 | 175256 | A | G | intergenic_region | MODIFIER | KCND2-KCND2 |
| NW_006532398.1 | 200122 | C | T | intergenic_region | MODIFIER | KCND2-KCND2 |
| NW_006532398.1 | 255146 | G | C | intergenic_region | MODIFIER | KCND2-KCND2 |
| NW_006532398.1 | 302035 | C | A | intergenic_region | MODIFIER | KCND2-KCND2 |
| NW_006532550.1 | 14469 | G | A | intergenic_region | MODIFIER | |
| NW_006532550.1 | 230463 | G | A | intergenic_region | MODIFIER | |
| NW_006532550.1 | 277473 | C | G | intergenic_region | MODIFIER | |
| NW_006532550.1 | 318564 | T | A,G | intergenic_region | MODIFIER | |
| NW_006532584.1 | 80530 | T | A | intergenic_region | MODIFIER | ZNF800-LOC103062773 |
| NW_006532656.1 | 47916 | G | C | intron_variant | MODIFIER | LOC112540401 |
| NW_006532656.1 | 49564 | A | G | intron_variant | MODIFIER | LOC112540401 |
| NW_006532656.1 | 124435 | G | A | intron_variant | MODIFIER | NRCAM |
| NW_006532656.1 | 125260 | C | T | intron_variant | MODIFIER | NRCAM |
| NW_006532656.1 | 128790 | C | A | intron_variant | MODIFIER | NRCAM |
| NW_006532656.1 | 158898 | G | T | downstream_gene_variant | MODIFIER | NRCAM |
| NW_006532656.1 | 164981 | A | G | intergenic_region | MODIFIER | NRCAM-CNTN1 |
| NW_006532656.1 | 172050 | A | G | upstream_gene_variant | MODIFIER | CNTN1 |
| NW_006532656.1 | 229223 | T | C | intron_variant | MODIFIER | CNTN1 |
| NW_006532656.1 | 238348 | G | A | intron_variant | MODIFIER | CNTN1 |
| NW_006532656.1 | 246036 | T | A | intron_variant | MODIFIER | CNTN1 |
| NW_006532656.1 | 289636 | C | T | intron_variant | MODIFIER | CNTN1 |
| NW_006532656.1 | 303084 | A | G | intron_variant | MODIFIER | CNTN1 |
| NW_006532656.1 | 335942 | G | A | 3_prime_UTR_variant | MODIFIER | CNTN1 |

| | | | | | | |
|---|---|---|---|---|---|---|
| NW_006532734.1 | 88252 | A | T | intergenic_region | MODIFIER | GXYLT1-PDZRN4 |
| NW_006532734.1 | 281387 | C | G | intron_variant | MODIFIER | PDZRN4 |
| NW_006532762.1 | 6396 | C | T | intergenic_region | MODIFIER | CHR_START-ZNF277 |
| NW_006532762.1 | 25741 | C | T | intergenic_region | MODIFIER | CHR_START-ZNF277 |
| NW_006532762.1 | 32304 | T | C | intergenic_region | MODIFIER | CHR_START-ZNF277 |
| NW_006532762.1 | 46234 | G | C | intron_variant | MODIFIER | ZNF277 |
| NW_006532762.1 | 55450 | G | C,T | XM_007426683.3 | protein_coding | ZNF277 |
| NW_006532762.1 | 78013 | C | G | upstream_gene_variant | MODIFIER | ZNF277 |
| NW_006532762.1 | 87278 | A | G | intron_variant | MODIFIER | DOCK4 |
| NW_006532762.1 | 91115 | G | A | intron_variant | MODIFIER | DOCK4 |
| NW_006532762.1 | 111863 | C | T | intron_variant | MODIFIER | DOCK4 |
| NW_006532762.1 | 120413 | T | G | intron_variant | MODIFIER | DOCK4 |
| NW_006532762.1 | 135064 | C | T | intron_variant | MODIFIER | DOCK4 |
| NW_006532762.1 | 144499 | A | T | intron_variant | MODIFIER | DOCK4 |
| NW_006532762.1 | 144985 | C | T | intron_variant | MODIFIER | DOCK4 |
| NW_006532762.1 | 150716 | G | T | intron_variant | MODIFIER | DOCK4 |
| NW_006532762.1 | 176421 | A | G | intron_variant | MODIFIER | DOCK4 |
| NW_006532762.1 | 176740 | G | T | intron_variant | MODIFIER | DOCK4 |
| NW_006532762.1 | 177277 | G | A | intron_variant | MODIFIER | DOCK4 |
| NW_006532762.1 | 177501 | G | A | intron_variant | MODIFIER | DOCK4 |
| NW_006532762.1 | 177905 | G | A | intron_variant | MODIFIER | DOCK4 |
| NW_006532762.1 | 178255 | T | C | intron_variant | MODIFIER | DOCK4 |
| NW_006532762.1 | 178470 | C | T | intron_variant | MODIFIER | DOCK4 |
| NW_006532762.1 | 178781 | T | G | intron_variant | MODIFIER | DOCK4 |
| NW_006532762.1 | 179065 | G | A | intron_variant | MODIFIER | DOCK4 |
| NW_006532762.1 | 179330 | C | T | intron_variant | MODIFIER | DOCK4 |
| NW_006532762.1 | 181300 | C | T | intron_variant | MODIFIER | DOCK4 |
| NW_006532762.1 | 186955 | G | T | intron_variant | MODIFIER | DOCK4 |
| NW_006532762.1 | 187338 | A | G | intron_variant | MODIFIER | DOCK4 |
| NW_006532762.1 | 187503 | A | G | intron_variant | MODIFIER | DOCK4 |
| NW_006532762.1 | 187564 | C | T | intron_variant | MODIFIER | DOCK4 |

| | | | | | | | |
|---|---|---|---|---|---|---|---|
| NW_006532762.1 | 187771 | A | G | intron_variant | MODIFIER | DOCK4 | |
| NW_006532762.1 | 188373 | A | G | intron_variant | MODIFIER | DOCK4 | |
| NW_006532762.1 | 189689 | C | T | intron_variant | MODIFIER | DOCK4 | |
| NW_006532762.1 | 190352 | G | A | intron_variant | MODIFIER | DOCK4 | |
| NW_006532762.1 | 190475 | A | G | intron_variant | MODIFIER | DOCK4 | |
| NW_006532762.1 | 192104 | A | G | intron_variant | MODIFIER | DOCK4 | |
| NW_006532762.1 | 193300 | C | T | intron_variant | MODIFIER | DOCK4 | |
| NW_006532762.1 | 210368 | A | G | intron_variant | MODIFIER | DOCK4 | |
| NW_006532762.1 | 255860 | G | A | intron_variant | MODIFIER | DOCK4 | |
| NW_006532762.1 | 259975 | T | A | intron_variant | MODIFIER | DOCK4 | |
| NW_006532762.1 | 291620 | A | G | intron_variant | MODIFIER | DOCK4 | |
| NW_006532992.1 | 255325 | C | G | 5_prime_UTR_variant | MODIFIER | ALG12 | |
| NW_006533029.1 | 6680 | G | C | intergenic_region | MODIFIER | CHR_START-FOXP2 | |
| NW_006533029.1 | 13107 | G | C | intergenic_region | MODIFIER | CHR_START-FOXP2 | |
| NW_006533029.1 | 70500 | G | C | intron_variant | MODIFIER | FOXP2 | |
| NW_006533029.1 | 98314 | C | T | intron_variant | MODIFIER | FOXP2 | |
| NW_006533029.1 | 118917 | C | T | intron_variant | MODIFIER | FOXP2 | |
| NW_006533029.1 | 119689 | A | C | intron_variant | MODIFIER | FOXP2 | |
| NW_006533029.1 | 126526 | G | C | intron_variant | MODIFIER | FOXP2 | |
| NW_006533029.1 | 166213 | A | T | intron_variant | MODIFIER | FOXP2 | |
| NW_006533029.1 | 168099 | A | G | intron_variant | MODIFIER | FOXP2 | |
| NW_006533029.1 | 170111 | G | T,C | XM_025166700.1 | protein_coding | | 17-Mar |
| NW_006533029.1 | 178330 | G | A | intron_variant | MODIFIER | FOXP2 | |
| NW_006533029.1 | 182255 | T | C | intron_variant | MODIFIER | FOXP2 | |
| NW_006533029.1 | 190096 | A | G | intron_variant | MODIFIER | FOXP2 | |
| NW_006533029.1 | 191513 | G | A | intron_variant | MODIFIER | FOXP2 | |
| NW_006533029.1 | 229787 | T | C | intron_variant | MODIFIER | FOXP2 | |
| NW_006533029.1 | 248375 | G | A | intron_variant | MODIFIER | FOXP2 | |
| NW_006533029.1 | 251642 | T | A | intron_variant | MODIFIER | FOXP2 | |
| NW_006533029.1 | 265610 | G | A | intron_variant | MODIFIER | FOXP2 | |
| NW_006533029.1 | 266802 | C | T | intron_variant | MODIFIER | FOXP2 | |

| | | | | | | |
|---|---|---|---|---|---|---|
| NW_006533029.1 | 288048 | A | G | intergenic_region | MODIFIER | FOXP2-CHR_END |
| NW_006533029.1 | 304541 | C | T | intergenic_region | MODIFIER | FOXP2-CHR_END |
| NW_006533320.1 | 58210 | A | G | intron_variant | MODIFIER | MEIG1 |
| NW_006533414.1 | 4481 | T | C | intergenic_region | MODIFIER | CHR_START-SMIM30 |
| NW_006533414.1 | 5664 | T | G,A | CHR_START-SMIM30 | | |
| NW_006533414.1 | 33268 | G | A | intergenic_region | MODIFIER | CHR_START-SMIM30 |
| NW_006533414.1 | 41039 | A | G | intergenic_region | MODIFIER | CHR_START-SMIM30 |
| NW_006533414.1 | 46458 | C | T | intergenic_region | MODIFIER | CHR_START-SMIM30 |
| NW_006533414.1 | 52330 | C | T | intergenic_region | MODIFIER | CHR_START-SMIM30 |
| NW_006533414.1 | 52985 | A | T | intergenic_region | MODIFIER | CHR_START-SMIM30 |
| NW_006533414.1 | 53295 | C | CTT | intergenic_region | MODIFIER | CHR_START-SMIM30 |
| NW_006533414.1 | 60379 | A | G | intergenic_region | MODIFIER | CHR_START-SMIM30 |
| NW_006533414.1 | 63980 | T | C | intergenic_region | MODIFIER | CHR_START-SMIM30 |
| NW_006533414.1 | 68613 | C | G | intergenic_region | MODIFIER | CHR_START-SMIM30 |
| NW_006533414.1 | 68905 | T | G | intergenic_region | MODIFIER | CHR_START-SMIM30 |
| NW_006533414.1 | 69074 | G | T | intergenic_region | MODIFIER | CHR_START-SMIM30 |
| NW_006533414.1 | 84339 | T | C | intergenic_region | MODIFIER | CHR_START-SMIM30 |
| NW_006533414.1 | 84828 | T | A | intergenic_region | MODIFIER | CHR_START-SMIM30 |
| NW_006533414.1 | 89082 | T | C | intergenic_region | MODIFIER | CHR_START-SMIM30 |
| NW_006533414.1 | 90049 | G | C | intergenic_region | MODIFIER | CHR_START-SMIM30 |
| NW_006533414.1 | 93710 | C | T | intergenic_region | MODIFIER | CHR_START-SMIM30 |
| NW_006533414.1 | 93787 | G | C | intergenic_region | MODIFIER | CHR_START-SMIM30 |
| NW_006533414.1 | 93931 | G | A | intergenic_region | MODIFIER | CHR_START-SMIM30 |
| NW_006533414.1 | 94555 | A | G | intergenic_region | MODIFIER | CHR_START-SMIM30 |
| NW_006533414.1 | 95316 | G | A | intergenic_region | MODIFIER | CHR_START-SMIM30 |
| NW_006533414.1 | 95484 | G | C | intergenic_region | MODIFIER | CHR_START-SMIM30 |
| NW_006533414.1 | 97761 | G | A | intergenic_region | MODIFIER | CHR_START-SMIM30 |
| NW_006533414.1 | 98825 | C | T | intergenic_region | MODIFIER | CHR_START-SMIM30 |
| NW_006533414.1 | 99408 | A | C | intergenic_region | MODIFIER | CHR_START-SMIM30 |
| NW_006533414.1 | 101008 | G | A | intergenic_region | MODIFIER | CHR_START-SMIM30 |
| NW_006533414.1 | 101331 | A | G | intergenic_region | MODIFIER | CHR_START-SMIM30 |

| | | | | | | |
|---|---|---|---|---|---|---|
| NW_006533414.1 | 101875 | C | T | intergenic_region | MODIFIER | CHR_START-SMIM30 |
| NW_006533414.1 | 102752 | T | C | intergenic_region | MODIFIER | CHR_START-SMIM30 |
| NW_006533414.1 | 110672 | G | A | upstream_gene_variant | MODIFIER | SMIM30 |
| NW_006533414.1 | 118187 | A | G | downstream_gene_variant | MODIFIER | SMIM30 |
| NW_006533414.1 | 139165 | C | T | intron_variant | MODIFIER | GPR85 |
| NW_006533414.1 | 144924 | C | G | downstream_gene_variant | MODIFIER | GPR85 |
| NW_006533414.1 | 157613 | G | A | intergenic_region | MODIFIER | GPR85-LOC112540885 |
| NW_006533414.1 | 162097 | A | G | intergenic_region | MODIFIER | GPR85-LOC112540885 |
| NW_006533414.1 | 174191 | C | G | intergenic_region | MODIFIER | GPR85-LOC112540885 |
| NW_006533414.1 | 206553 | C | G | upstream_gene_variant | MODIFIER | BMT2 |
| NW_006533414.1 | 221760 | C | G | 3_prime_UTR_variant | MODIFIER | BMT2 |
| NW_006533414.1 | 221985 | T | A | 3_prime_UTR_variant | MODIFIER | BMT2 |
| NW_006533704.1 | 22326 | A | G | intergenic_region | MODIFIER | |
| NW_006533704.1 | 87436 | T | C | intergenic_region | MODIFIER | |
| NW_006533704.1 | 87543 | A | T | intergenic_region | MODIFIER | |
| NW_006533704.1 | 108677 | C | G | intergenic_region | MODIFIER | |
| NW_006533704.1 | 135119 | G | A | intergenic_region | MODIFIER | |
| NW_006533704.1 | 137455 | G | C | intergenic_region | MODIFIER | |
| NW_006533704.1 | 139320 | T | C | intergenic_region | MODIFIER | |
| NW_006533704.1 | 149143 | T | C | intergenic_region | MODIFIER | |
| NW_006533704.1 | 174318 | G | A | intergenic_region | MODIFIER | |
| NW_006533704.1 | 189697 | T | C | intergenic_region | MODIFIER | |
| NW_006533704.1 | 195596 | A | G | intergenic_region | MODIFIER | |
| NW_006533753.1 | 1353 | A | C,G | CHR_START-MDFIC | | |
| NW_006533753.1 | 15450 | T | C | intergenic_region | MODIFIER | CHR_START-MDFIC |
| NW_006533753.1 | 41106 | T | C | intergenic_region | MODIFIER | CHR_START-MDFIC |
| NW_006533753.1 | 45951 | C | T | intergenic_region | MODIFIER | CHR_START-MDFIC |
| NW_006533753.1 | 51544 | C | A | intergenic_region | MODIFIER | CHR_START-MDFIC |
| NW_006533753.1 | 61807 | G | C | intergenic_region | MODIFIER | CHR_START-MDFIC |
| NW_006533753.1 | 62335 | T | C | intergenic_region | MODIFIER | CHR_START-MDFIC |
| NW_006533753.1 | 100969 | G | A | intergenic_region | MODIFIER | CHR_START-MDFIC |

| | | | | | | |
|---|---|---|---|---|---|---|
| NW_006533753.1 | 109750 | C | T | intergenic_region | MODIFIER | CHR_START-MDFIC |
| NW_006533753.1 | 119078 | C | T | intergenic_region | MODIFIER | CHR_START-MDFIC |
| NW_006533753.1 | 120752 | A | C | intergenic_region | MODIFIER | CHR_START-MDFIC |
| NW_006533753.1 | 131834 | A | G,T | CHR_START-MDFIC | | |
| NW_006533753.1 | 187318 | A | G | intergenic_region | MODIFIER | CHR_START-MDFIC |
| NW_006533753.1 | 202488 | A | C | intergenic_region | MODIFIER | CHR_START-MDFIC |
| NW_006533753.1 | 207762 | C | T | intergenic_region | MODIFIER | CHR_START-MDFIC |
| NW_006533850.1 | 18242 | C | T | intron_variant | MODIFIER | LOC103067393 |
| NW_006533850.1 | 60029 | C | T | intergenic_region | MODIFIER | LOC103067393-MET |
| NW_006533850.1 | 62912 | C | G | intergenic_region | MODIFIER | LOC103067393-MET |
| NW_006533850.1 | 105238 | G | A | intergenic_region | MODIFIER | LOC103067393-MET |
| NW_006533850.1 | 111276 | C | T | 5_prime_UTR_variant | MODIFIER | MET |
| NW_006533850.1 | 220015 | C | T | downstream_gene_variant | MODIFIER | MET |
| NW_006534020.1 | 11503 | G | C | intron_variant | MODIFIER | TES |
| NW_006534020.1 | 14140 | C | T | intron_variant | MODIFIER | TES |
| NW_006534020.1 | 25295 | C | T | intron_variant | MODIFIER | TES |
| NW_006534020.1 | 39582 | T | A | upstream_gene_variant | MODIFIER | TES |
| NW_006534020.1 | 57253 | T | C | intron_variant | MODIFIER | TFEC |
| NW_006534020.1 | 74156 | A | C | intron_variant | MODIFIER | TFEC |
| NW_006534020.1 | 78482 | C | T | intron_variant | MODIFIER | TFEC |
| NW_006534020.1 | 78923 | A | C | intron_variant | MODIFIER | TFEC |
| NW_006534020.1 | 81206 | T | C | intron_variant | MODIFIER | TFEC |
| NW_006534020.1 | 84289 | A | G | intron_variant | MODIFIER | TFEC |
| NW_006534020.1 | 84462 | G | T | intron_variant | MODIFIER | TFEC |
| NW_006534020.1 | 86561 | C | T | intron_variant | MODIFIER | TFEC |
| NW_006534020.1 | 89179 | C | T | intron_variant | MODIFIER | TFEC |
| NW_006534020.1 | 90897 | A | G | intron_variant | MODIFIER | TFEC |
| NW_006534020.1 | 93724 | T | G | intron_variant | MODIFIER | TFEC |
| NW_006534020.1 | 94369 | C | G | intron_variant | MODIFIER | TFEC |
| NW_006534020.1 | 94707 | T | C | intron_variant | MODIFIER | TFEC |
| NW_006534020.1 | 97367 | G | A | intron_variant | MODIFIER | TFEC |

| | | | | | | |
|---|---|---|---|---|---|---|
| NW_006534020.1 | 98473 | C | T | intron_variant | MODIFIER | TFEC |
| NW_006534020.1 | 100782 | C | T | intron_variant | MODIFIER | TFEC |
| NW_006534020.1 | 106194 | A | C | intron_variant | MODIFIER | TFEC |
| NW_006534020.1 | 107938 | C | T | intron_variant | MODIFIER | TFEC |
| NW_006534020.1 | 115640 | A | G | intron_variant | MODIFIER | TFEC |
| NW_006534020.1 | 117555 | G | A | intron_variant | MODIFIER | TFEC |
| NW_006534020.1 | 118714 | C | T | intron_variant | MODIFIER | TFEC |
| NW_006534020.1 | 122527 | C | T | intron_variant | MODIFIER | TFEC |
| NW_006534020.1 | 128782 | T | C | intron_variant | MODIFIER | TFEC |
| NW_006534020.1 | 144397 | T | C | intron_variant | MODIFIER | TFEC |
| NW_006534020.1 | 145047 | G | A | intron_variant | MODIFIER | TFEC |
| NW_006534020.1 | 145788 | T | C | intron_variant | MODIFIER | TFEC |
| NW_006534020.1 | 152557 | T | A | intron_variant | MODIFIER | TFEC |
| NW_006534020.1 | 160458 | C | T | stop_gained | HIGH | TFEC |
| NW_006534020.1 | 161049 | A | G | intron_variant | MODIFIER | TFEC |
| NW_006534020.1 | 172658 | G | A | downstream_gene_variant | MODIFIER | TFEC |
| NW_006534020.1 | 174250 | C | T | downstream_gene_variant | MODIFIER | TFEC |
| NW_006534020.1 | 177118 | G | A | intergenic_region | MODIFIER | TFEC-CHR_END |
| NW_006534020.1 | 184905 | A | G | intergenic_region | MODIFIER | TFEC-CHR_END |
| NW_006534020.1 | 187003 | C | T | intergenic_region | MODIFIER | TFEC-CHR_END |
| NW_006534020.1 | 189636 | C | G | intergenic_region | MODIFIER | TFEC-CHR_END |
| NW_006534020.1 | 190461 | C | A | intergenic_region | MODIFIER | TFEC-CHR_END |
| NW_006534020.1 | 190480 | A | G | intergenic_region | MODIFIER | TFEC-CHR_END |
| NW_006534020.1 | 195021 | T | C | intergenic_region | MODIFIER | TFEC-CHR_END |
| NW_006534020.1 | 204352 | C | T | intergenic_region | MODIFIER | TFEC-CHR_END |
| NW_006534273.1 | 75867 | G | A | intergenic_region | MODIFIER | |
| NW_006534273.1 | 115761 | G | A | intergenic_region | MODIFIER | |
| NW_006534273.1 | 172918 | T | C | intergenic_region | MODIFIER | |
| NW_006534273.1 | 190897 | C | T | intergenic_region | MODIFIER | |
| NW_006534273.1 | 195145 | A | G | intergenic_region | MODIFIER | |
| NW_006534432.1 | 4777 | C | T | intergenic_region | MODIFIER | CHR_START-PPP1R3A |

| | | | | | | |
|---|---|---|---|---|---|---|
| NW_006534432.1 | 7225 | T | C | intergenic_region | MODIFIER | CHR_START-PPP1R3A |
| NW_006534432.1 | 30900 | T | A | intergenic_region | MODIFIER | CHR_START-PPP1R3A |
| NW_006534432.1 | 38162 | A | G | intergenic_region | MODIFIER | CHR_START-PPP1R3A |
| NW_006534432.1 | 42369 | A | C | downstream_gene_variant | MODIFIER | PPP1R3A |
| NW_006534432.1 | 69206 | A | G | intron_variant | MODIFIER | PPP1R3A |
| NW_006534432.1 | 75363 | C | A | upstream_gene_variant | MODIFIER | PPP1R3A |
| NW_006534432.1 | 82810 | G | A | intergenic_region | MODIFIER | PPP1R3A-CHR_END |
| NW_006534432.1 | 85655 | T | G | intergenic_region | MODIFIER | PPP1R3A-CHR_END |
| NW_006534432.1 | 89156 | G | A | intergenic_region | MODIFIER | PPP1R3A-CHR_END |
| NW_006534432.1 | 93102 | C | A | intergenic_region | MODIFIER | PPP1R3A-CHR_END |
| NW_006534432.1 | 96774 | A | G | intergenic_region | MODIFIER | PPP1R3A-CHR_END |
| NW_006534432.1 | 108409 | T | C | intergenic_region | MODIFIER | PPP1R3A-CHR_END |
| NW_006534432.1 | 114574 | A | G | intergenic_region | MODIFIER | PPP1R3A-CHR_END |
| NW_006534432.1 | 114887 | G | A | intergenic_region | MODIFIER | PPP1R3A-CHR_END |
| NW_006534432.1 | 137974 | G | C | intergenic_region | MODIFIER | PPP1R3A-CHR_END |
| NW_006534432.1 | 154471 | C | G | intergenic_region | MODIFIER | PPP1R3A-CHR_END |
| NW_006534489.1 | 12880 | C | A | intergenic_region | MODIFIER | |
| NW_006534489.1 | 19335 | G | C | intergenic_region | MODIFIER | |
| NW_006534489.1 | 19508 | G | A | intergenic_region | MODIFIER | |
| NW_006534489.1 | 51739 | C | T | intergenic_region | MODIFIER | |
| NW_006534489.1 | 90237 | G | A | intergenic_region | MODIFIER | |
| NW_006534489.1 | 130225 | G | C | intergenic_region | MODIFIER | |
| NW_006535328.1 | 114291 | T | A | intergenic_region | MODIFIER | LOC103055319-LOC103055319 |
| NW_006535573.1 | 8749 | C | T | intron_variant | MODIFIER | CAPZA2 |
| NW_006535573.1 | 9302 | G | C | intron_variant | MODIFIER | CAPZA2 |
| NW_006535573.1 | 11018 | A | T | intron_variant | MODIFIER | CAPZA2 |
| NW_006535573.1 | 11286 | G | A | intron_variant | MODIFIER | CAPZA2 |
| NW_006535573.1 | 11513 | A | T | intron_variant | MODIFIER | CAPZA2 |
| NW_006535573.1 | 13629 | C | T | intron_variant | MODIFIER | CAPZA2 |
| NW_006535573.1 | 16200 | A | T | intron_variant | MODIFIER | CAPZA2 |

| | | | | | | | |
|---|---|---|---|---|---|---|---|
| NW_006535573.1 | 16955 | A | T | intron_variant | MODIFIER | CAPZA2 | |
| NW_006535573.1 | 19096 | G | A | intron_variant | MODIFIER | CAPZA2 | |
| NW_006535573.1 | 21140 | G | A | intron_variant | MODIFIER | CAPZA2 | |
| NW_006535573.1 | 23728 | G | A | intron_variant | MODIFIER | CAPZA2 | |
| NW_006535573.1 | 24989 | C | G | intron_variant | MODIFIER | CAPZA2 | |
| NW_006535573.1 | 25767 | G | A | intron_variant | MODIFIER | CAPZA2 | |
| NW_006535573.1 | 27051 | C | G | intron_variant | MODIFIER | CAPZA2 | |
| NW_006535573.1 | 30210 | G | A,T | XM_007437135.3 | protein_coding | | 10-Oct |
| NW_006535573.1 | 54839 | T | C | upstream_gene_variant | MODIFIER | ST7 | |
| NW_006535573.1 | 126162 | T | C | intron_variant | MODIFIER | ST7 | |
| NW_006535573.1 | 136157 | T | G | intron_variant | MODIFIER | ST7 | |
| NW_006535650.1 | 9082 | C | T | intergenic_region | MODIFIER | CHR_START-LOC103055238 | |
| NW_006535650.1 | 33998 | G | A | intergenic_region | MODIFIER | CHR_START-LOC103055238 | |
| NW_006535650.1 | 37546 | G | A | intergenic_region | MODIFIER | CHR_START-LOC103055238 | |
| NW_006535650.1 | 61741 | G | A | intergenic_region | MODIFIER | CHR_START-LOC103055238 | |
| NW_006535650.1 | 71564 | C | T | intergenic_region | MODIFIER | CHR_START-LOC103055238 | |
| NW_006535650.1 | 72082 | G | A | intergenic_region | MODIFIER | CHR_START-LOC103055238 | |
| NW_006535650.1 | 128989 | C | T | intergenic_region | MODIFIER | LOC103055238-CHR_END | |
| NW_006535793.1 | 48357 | G | A | intergenic_region | MODIFIER | | |
| NW_006536153.1 | 10186 | A | C | intergenic_region | MODIFIER | CHR_START-FAM3C | |
| NW_006536169.1 | 31331 | G | A | intergenic_region | MODIFIER | | |
| NW_006536169.1 | 71768 | G | C | intergenic_region | MODIFIER | | |
| NW_006536169.1 | 94565 | A | C | intergenic_region | MODIFIER | | |
| NW_006536169.1 | 114527 | C | T | intergenic_region | MODIFIER | | |
| NW_006536323.1 | 27429 | A | G | intergenic_region | MODIFIER | LOC103066484-LOC112541966 | |
| NW_006536323.1 | 40910 | A | G | intergenic_region | MODIFIER | LOC112541966-CHR_END | |
| NW_006536993.1 | 18595 | C | A | intergenic_region | MODIFIER | | |
| NW_006536993.1 | 35881 | C | G | intergenic_region | MODIFIER | | |
| NW_006536993.1 | 75875 | G | C | intergenic_region | MODIFIER | | |
| NW_006537005.1 | 71370 | C | T | intergenic_region | MODIFIER | | |

| | | | | | |
|---|---|---|---|---|---|
| NW_006537198.1 | 29734 | T | A | intergenic_region | MODIFIER |
| NW_006537198.1 | 42905 | T | C | intergenic_region | MODIFIER |
| NW_006537198.1 | 49563 | C | T | intergenic_region | MODIFIER |
| NW_006537198.1 | 53483 | C | G | intergenic_region | MODIFIER |
| NW_006537198.1 | 57934 | T | G | intergenic_region | MODIFIER |
| NW_006537198.1 | 57958 | T | G | intergenic_region | MODIFIER |
| NW_006537198.1 | 61528 | G | A | intergenic_region | MODIFIER |
| NW_006537198.1 | 70698 | T | C | intergenic_region | MODIFIER |
| NW_006537198.1 | 81626 | C | T | intergenic_region | MODIFIER |
| NW_006537198.1 | 82903 | T | C | intergenic_region | MODIFIER |
| NW_006537198.1 | 83679 | G | T | intergenic_region | MODIFIER |
| NW_006537513.1 | 3569 | A | G | intergenic_region | MODIFIER |
| NW_006537513.1 | 6143 | A | G | intergenic_region | MODIFIER |
| NW_006537513.1 | 13553 | C | T | intergenic_region | MODIFIER |
| NW_006537513.1 | 26446 | C | A | intergenic_region | MODIFIER |
| NW_006537513.1 | 50353 | T | C | intergenic_region | MODIFIER |
| NW_006537513.1 | 58581 | A | G | intergenic_region | MODIFIER |
| NW_006537513.1 | 69339 | A | C | intergenic_region | MODIFIER |
| NW_006537513.1 | 70842 | T | A | intergenic_region | MODIFIER |
| NW_006538027.1 | 4234 | C | T | intergenic_region | MODIFIER |
| NW_006538027.1 | 7719 | A | T | intergenic_region | MODIFIER |
| NW_006538027.1 | 17519 | C | G | intergenic_region | MODIFIER |
| NW_006538027.1 | 19802 | C | G | intergenic_region | MODIFIER |
| NW_006538027.1 | 26667 | A | T | intergenic_region | MODIFIER |
| NW_006538027.1 | 31092 | A | C | intergenic_region | MODIFIER |
| NW_006538027.1 | 35496 | G | A | intergenic_region | MODIFIER |
| NW_006538239.1 | 14753 | A | G | intergenic_region | MODIFIER |
| NW_006538239.1 | 20125 | C | T | intergenic_region | MODIFIER |
| NW_006538239.1 | 21067 | C | T | intergenic_region | MODIFIER |
| NW_006538239.1 | 23109 | G | C | intergenic_region | MODIFIER |
| NW_006538239.1 | 26846 | C | G | intergenic_region | MODIFIER |

| | | | | | | |
|---|---|---|---|---|---|---|
| NW_006538239.1 | 36729 | G | A | intergenic_region | MODIFIER | |
| NW_006538239.1 | 61058 | G | A | intergenic_region | MODIFIER | |
| NW_006538428.1 | 16356 | C | G | intergenic_region | MODIFIER | |
| NW_006538570.1 | 10922 | T | C | intergenic_region | MODIFIER | |
| NW_006538570.1 | 17178 | T | G | intergenic_region | MODIFIER | |
| NW_006538570.1 | 20843 | T | C | intergenic_region | MODIFIER | |
| NW_006538570.1 | 24004 | T | C | intergenic_region | MODIFIER | |
| NW_006538570.1 | 26373 | G | C | intergenic_region | MODIFIER | |
| NW_006538570.1 | 26509 | G | A | intergenic_region | MODIFIER | |
| NW_006538570.1 | 27509 | A | G | intergenic_region | MODIFIER | |
| NW_006538570.1 | 29132 | T | C | intergenic_region | MODIFIER | |
| NW_006538570.1 | 29248 | G | A | intergenic_region | MODIFIER | |
| NW_006538666.1 | 50308 | C | T | intergenic_region | MODIFIER | |
| NW_006538666.1 | 54558 | T | G | intergenic_region | MODIFIER | |
| NW_006538920.1 | 29179 | G | T | intergenic_region | MODIFIER | |
| NW_006538963.1 | 43657 | T | C | downstream_gene_variant | MODIFIER | LOC103064888 |
| NW_006539084.1 | 21291 | C | T | downstream_gene_variant | MODIFIER | IFRD1 |
| NW_006539084.1 | 22623 | C | G | intron_variant | MODIFIER | LSMEM1 |
| NW_006539084.1 | 22636 | C | T | intron_variant | MODIFIER | LSMEM1 |
| NW_006539084.1 | 23347 | C | T | intron_variant | MODIFIER | LSMEM1 |
| NW_006539084.1 | 26867 | C | T | synonymous_variant | LOW | LSMEM1 |
| NW_006539084.1 | 28392 | A | T | intron_variant | MODIFIER | LSMEM1 |
| NW_006539084.1 | 28792 | A | G | intron_variant | MODIFIER | LSMEM1 |
| NW_006539084.1 | 29041 | G | A | intron_variant | MODIFIER | LSMEM1 |
| NW_006539084.1 | 29785 | G | C | intron_variant | MODIFIER | LSMEM1 |
| NW_006539084.1 | 31618 | C | T | 3_prime_UTR_variant | MODIFIER | LSMEM1 |
| NW_006539084.1 | 31899 | G | A | 3_prime_UTR_variant | MODIFIER | LSMEM1 |
| NW_006539084.1 | 31923 | G | C | 3_prime_UTR_variant | MODIFIER | LSMEM1 |
| NW_006539084.1 | 42931 | A | G | intron_variant | MODIFIER | LOC112542615 |
| NW_006540138.1 | 22512 | A | C | intergenic_region | MODIFIER | |
| NW_006540138.1 | 25311 | G | A | intergenic_region | MODIFIER | |

| | | | | | | |
|---|---|---|---|---|---|---|
| NW_006541234.1 | 4988 | C | T | intergenic_region | MODIFIER | CHR_START-LRRN3 |
| NW_006541234.1 | 9596 | A | G | downstream_gene_variant | MODIFIER | LRRN3 |
| NW_006541234.1 | 11393 | C | G | downstream_gene_variant | MODIFIER | LRRN3 |
| NW_006542015.1 | 3980 | C | G | intergenic_region | MODIFIER | |
| NW_006542817.1 | 5157 | C | G | intergenic_region | MODIFIER | |
| NW_006544037.1 | 5429 | T | C | intergenic_region | MODIFIER | |
| NW_006544037.1 | 5566 | G | A | intergenic_region | MODIFIER | |
| NW_006545079.1 | 1022 | C | T | intergenic_region | MODIFIER | |
| NW_006545079.1 | 2021 | T | C | intergenic_region | MODIFIER | |
| NW_006570921.1 | 206 | A | T | intergenic_region | MODIFIER | |
